# SOX2 expression identifies Ewing sarcoma patients with high risk for tumor relapse and poor survival

**DOI:** 10.1101/498253

**Authors:** G. Sannino, A. Marchetto, A. Ranft, S. Jabar, C. Zacherl, R. Alba-Rubio, S. Stein, F. S. Wehweck, M. M. Kiran, T. L. B. Hölting, J. Musa, L. Romero-Pérez, F. Cidre-Aranaz, M. M. L. Knott, J. Li, H. Jürgens, A. Sastre, J. Alonso, W. Da Silveira, G. Hardiman, J. S. Gerke, M. F. Orth, W. Hartmann, T. Kirchner, S. Ohmura, U. Dirksen, T. G. P. Grünewald

## Abstract

**Purpose:** Up to 30–40% of Ewing sarcoma (EwS) patients with non-metastatic disease develop local or metastatic relapse within a time range of 2–10 years. This is in part caused by the absence of prognostic biomarkers that can identify high-risk patients to assign them to risk-adapted monitoring and treatment regimens. Since cancer stemness has been associated with tumor relapse and poor outcome, we investigated in the current study the prognostic potential SOX2 (sex determining region Y box 2) – a major transcription factor involved in development and stemness – previously described to contribute to the undifferentiated phenotype of EwS.

**Methods:** Two independent patient cohorts, one consisting of 189 retrospectively collected EwS tumors with corresponding mRNA expression data (test cohort) and the other of 141 prospectively collected formalin-fixed and paraffin embedded resected tumors (validation cohort), were employed to analyze SOX2 expression levels by DNA microarrays or immunohistochemistry, respectively, and to compare them with clinical parameters and patient outcome. Two methods were employed to test the validity of the results at both mRNA and protein levels.

**Results:** Both cohorts showed that only a subset of EwS patients (16–20%) express high SOX2 mRNA or protein levels, which significantly correlated with poor overall survival. Multivariate analyses of our validation cohort revealed that high SOX2 expression represents a major risk-factor for survival (HR=3.19; *P*<0.01) that is independent from metastasis and other known clinical risk-factors at time of diagnosis. Univariate analysis demonstrated that SOX2-high expression correlated with tumor relapse (P=0.001). Median first relapse was at 14.7 months (range: 3.5–180.7).

**Conclusions:** High SOX2 expression constitutes an independent prognostic biomarker for EwS patients with poor outcome, which may help to identify patients with localized disease who are at high risk for tumor relapse within the first two years after diagnosis.

## Introduction

Ewing sarcoma (EwS) represents the second most common bone and soft tissue cancer in children and adolescents, which potentially arises from neuroectodermal or mesodermal mesenchymal stem cells^1,2^. In fact, EwS tumors display a largely undifferentiated and ‘stemness’ phenotype, which is believed to contribute to its clinical aggressiveness^1^. Genetically, the hallmark of EwS are chromosomal translocations generating chimeric proteins through fusion of the *EWSR1* (*Ewing sarcoma breakpoint region 1*) gene to variable members of the ETS (E26 transformation specific) family of transcription factors, in 85% *FLI1 (Friend leukemia virus integration 1*)^3–5^. These EWSR1-ETS fusion oncoproteins act as aberrant transcription factors that promote tumor initiation and progression by massively rewiring the cellular transcriptome and spliceosome^1,6,7^.

Sixty to 75% of EwS patients benefit from multimodal therapy^2,8^. Consequently, >30% of patients show limited response to treatment, which is often first noted based on assessment of histological response after neoadjuvant chemotherapy^9^. Since limited treatment response may be associated with early relapse^10^, upfront identification of high-risk patients is essential to assign them to adequate treatment regimens. Despite several clinicopathological features (such as tumor volume, histological response, tumor site and age at diagnosis) have been associated with high-risk disease, so far the presence of metastasis at diagnosis remains the major factor for risk-stratification in EwS patients^11^. However, for patients with localized disease, risk prediction remains difficult as no *bona fide* prognostic biomarkers independent from metastasis are available^1,12^.

A prior report indicated that stemness in EwS may be mediated via EWSR1-FLI-induced expression of *SOX2 (sex determining region Y box 2*)^13^ – a well-known stemness gene overexpressed in many cancers^14^. We therefore explored in the current study the expression pattern of SOX2 in two independent EwS cohorts and investigated whether it may serve as biomarker for outcome prediction in EwS.

We show that SOX2 is expressed only in ~16–20% of EwS tumors, which is associated with very poor patient outcome independent of metastasis. Moreover, high expression of SOX2 significantly correlates with tumor relapse. Our findings suggest that detection of high SOX2 expression by immunohistochemistry or RNA-based techniques may constitute a broadly available biomarker for risk-stratification even for patients with localized disease.

## MATERIALS AND METHODS

### Study populations

For this study, a retrospective test-cohort and a prospective validation-cohort were analyzed. Patients’ characteristics of both cohorts are listed in **Table 1**.

**Table 1.** Patient characteristics of the mRNA-cohort (*n*=189) and TMA-cohort (*n*=141). * all extremity localizations are non-axial; ** good response defined as <10% viable tumor cells; NA, not available.

| Variable | Test-cohort (mRNA)<br>*n* (%) | Validation-cohort (TMA)<br>*n* (%) |
| --- | --- | --- |
| Sex |  |  |
| male | 86 (57.7) | 87 (61.7) |
| female | 63 (42.3)<br>(missing: 47) | 54 (38.3) |
| Age at diagnosis |  |  |
| <15 years | 112 (59.3) | 70 (49.7) |
| ≥15 years | 77 (40.7) | 71 (50.3) |
| Metastasis at diagnosis |  |  |
| absent | 53 (69.7) | 87 (61.7) |
| present | 23 (30.3)<br>(missing: 113) | 54 (38.3) |
| Site* |  |  |
| axial | NA | 86 (61.9) |
| non-axial | NA | 53 (38.1)<br>(missing: 2) |
| Tumor volume |  |  |
| <200 ml | NA | 76 (55.9) |
| ≥200 ml | NA | 60 (44.1)<br>(missing: 5) |
| Histological response** |  |  |
| good | NA | 67 (79.8) |
| poor | NA | 17 (20.2)<br>(missing: 57) |
| Status |  |  |
| alive | 113 (59.8) | 84 (59.6) |
| dead | 76 (40.2) | 57 (40.4) |
| SOX2 expression |  |  |
| high | 38 (20.1) | 22 (15.6) |
| low | 151 (79.9) | 119 (84.4) |

The study population of the test-cohort consisted of 189 EwS patients whose molecularly confirmed and retrospectively collected primary tumors were profiled on the mRNA level by gene expression microarrays in previous studies^15–18^. To assemble this mRNA-cohort, microarray data generated on Affymetrix HG-U133Plus2.0, Affymetrix HuEx-1.0-st or Amersham/GE Healthcare CodeLink microarrays of 189 EwS tumors (Gene Expression Omnibus (GEO) accession codes: GSE63157^15^, GSE12102^16^, GSE17618^17^, GSE34620 ^18^, and unpublished data) provided with clinical annotations were normalized as previously described^19^. Batch effects were removed using the ComBat algorithm^20^.

The study population of the validation-cohort was composed of 141 EwS patients treated with first-line therapy according to the successive phase III Ewing sarcoma protocols of European Intergroup Cooperative Ewing’s Sarcoma Study (EICESS) 92 (1992 to 1998), and EURO-EWING 99 (1999 to 2009) run by the German Pediatric Society of Oncology and Hematology (GPOH). These studies were registered under ClinicalTrials.gov and approved by the appropriate ethics committees. The corresponding patient tumors were prospectively collected and tissue microarrays (TMAs) were constructed as detailed below. This study population included 87 males and 54 females.

### TMA establishment and immunohistochemistry (IHC)

Formalin-fixed paraffin-embedded (FFPE) EwS samples were retrieved from the archives of the Institute of Pathology of the LMU Munich (Germany) and the Gerhard-Domagk-Institute for Pathology of the University of Münster (Germany) with approval of the corresponding institutional review boards. All EwS samples were reviewed by a reference pathologist and diagnosis was confirmed either by detection of pathognomonic *EWSR1-ETS* fusion oncogenes by qRT-PCR or detection of an *EWSR1* break apart by fluorescence-in-situ-hybridization (FISH). TMAs were constructed as previously described^19^. Each EwS case was represented by at least two cores (each 1 mm in diameter).

For IHC, 4 μm sections were cut, and antigen retrieval was performed with microwave treatment. Slides were incubated with a primary monoclonal rabbit anti-SOX2-antibody (1:100 dilution, D6D9 XP Cell Signaling) for 60 min. Then slides were incubated with a secondary anti-rabbit IgG antibody (ImmPress Reagent Kit, Peroxidase-conjugated) followed by target detection using AECplus chromogen for 10 min (Dako, K3461). The specificity of the used anti-SOX2-antibody was confirmed in xenografts of the EwS cell line POE in NOD/Scid/gamma (NSG) mice, in which SOX2 expression was silenced by a doxycycline (dox-)inducible shRNA against the 3’UTR of *SOX2* (**Fig. S3B**). For quantification of SOX2 immunoreactivity, the average percentage of SOX2-positive nuclei was evaluated by a data-blinded pathologist by examining at least 5 high-power fields per case. Examples of nuclear SOX2 immunoreactivity are given in **Fig. 1E**.

**Figure 1:**
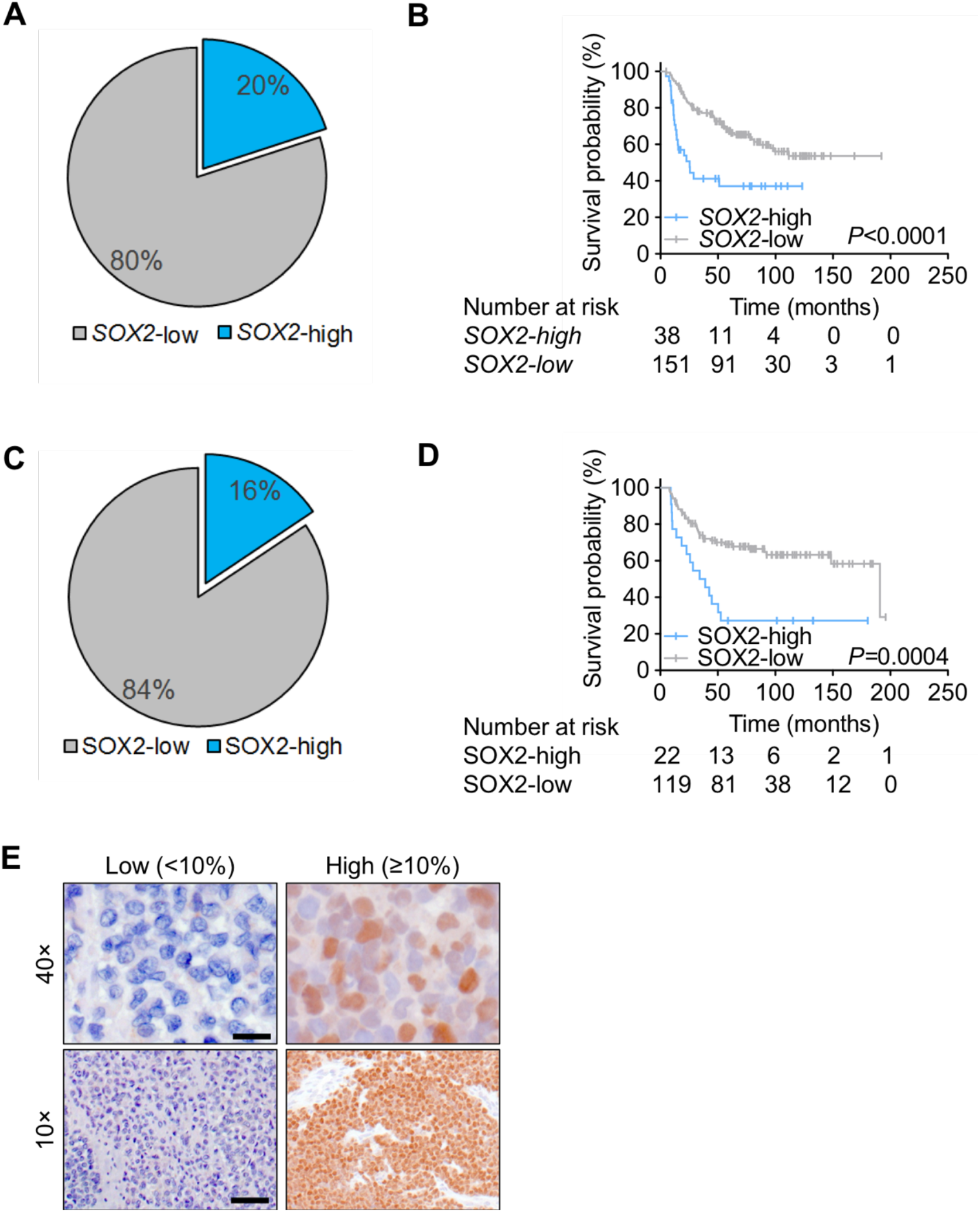
SOX2 is expressed in a subset of EwS patients and correlates with overall survival (OS) (A). Microarray analysis of SOX2 mRNA expression in 189 primary EwS cases. (B). Kaplan-Meier analysis of OS in 189 EwS patients stratified by their *SOX2* mRNA expression level (cut-off 80^th^ percentile). (C). Analysis of SOX2 protein expression levels in an independent cohort of 141 EwS patients in a TMA by IHC. (D). Kaplan-Meier analysis of OS in 141 EwS patients stratified by their SOX2 protein expression level (cut-off >10% positive nuclei). (E). Representative images for SOX2 IHC (nuclear staining) in EwS patients with different percentages of nuclear SOX2 immunoreactivity. Scale bars = 50 μM (40×) and 200 μM (10×).

### Statistical analyses

Statistical analyses were carried out with SPSS 19 (IBM Corporation, Armonk, NY) and SAS 9.2 (SAS Institute, Cary, NC) as described^21^. Overall survival (OS) was estimated by the Kaplan-Meier method. OS time was defined as the interval between the date of diagnosis and the date of last follow-up or death. Group comparisons were calculated by log-rank test. Multivariate analyses were carried out by applying the Cox proportional hazard method. Differences in proportions between groups were evaluated by chi-square or Fisher’s exact test. Significance level was set at *P*<0.05 for two-sided testing. No alpha corrections were carried out for multiple testing. Outcome was analyzed on an exploratory basis.

## RESULTS

### SOX2 is expressed in a subset of EwS patients and correlates with overall survival (OS)

SOX2 is a well-known transcription factor involved in stemness during normal development and cancer^14^. In 2010, Riggi *et al*. proposed that EWSR1-FLI1 confers stemness features to EwS cells via up-regulation of SOX2 ^13^. We therefore reasoned that SOX2 expression levels might be connected to patient outcome.

To test this hypothesis, a test cohort obtained from a large EwS gene expression dataset comprising 189 samples, for which matched clinical data were available (**Table 1**), was employed to evaluate mRNA levels of *SOX2*. In contrast to previous suggestions that SOX2 overexpression might constitute a general feature of EwS^13^, we found by microarray analysis that *SOX2* mRNA was not or only minimally expressed in 79.9% (151/189) of tumors. Indeed, only 20.1% (38/189) of samples exhibited moderate to strong *SOX2* expression levels (**Fig. 1A**). Stratifying our mRNA-cohort by a cut-off of the 80^th^ percentile of *SOX2* expression in *SOX2*-low or -high cases, we noted that patients with high intratumoral *SOX2* expression had a worse OS than patients with low *SOX2* expression (*P*<0.0001) (**Fig. 1B**).

To validate this finding on the protein level, we stained an independent validation TMA-cohort comprising 141 EwS cases with an anti-SOX2-antibody with proven specificity for EwS (**Fig. S3B**). Patient characteristics are reported in **Table 1**. Scoring of the percentage of nuclear SOX2 positive cells confirmed that the majority of EwS cases (84.4%, 119/141) showed ≤10% positive nuclei (classified as SOX2-low) while only a small subset of patients (15.6%, 22/141) had >10% positive nuclei (classified as SOX2-high) (**Fig. 1C,E**). Strikingly, applying the cut-off of >10% SOX2 positive nuclei fully confirmed the strong association of SOX2 expression with poor OS of EwS patients (*P*=0.0004) (**Fig. 1D**).

Collectively, these results demonstrate, for the first time, that high SOX2 expression is not a common feature of EwS, but that its high mRNA and protein levels may serve as a biomarker for outcome prediction.

### High SOX2 expression is a major risk-factor for tumor associated-death independent from metastasis in EwS

To identify factors that may influence patients’ prognosis, we performed a multivariate analysis in our validation TMA-cohort as here additional clinicopathological information beyond OS were available. Interestingly, multivariate analysis revealed that the major risk-factors were metastatic disease at M2 stage and SOX2-high expression with hazard ratios (HR) of 4.86 and 3.19 (both *P*<0.01), respectively. Instead, M0 (HR=R; *P*<0.01), M1 (*n*=24; HR=1.76, *P*=0.13), age (≥15 years, HR=1.34, *P*=0.28) and primary axial tumor site (HR=1.64, *P*=0.11) did not show a significant impact on survival (*n*=141; **Table 2**).

**Table 2.** Multivariate analysis in all patients of the TMA-cohort (*n*=141). Met, metastases. * reference (R).

| Variable | HR (95%CI) | *P* |
| --- | --- | --- |
| Risk group |  |  |
| M0 (no met; *n*=87) | R* | <0.01 |
| M1 (lung met; *n*=24) | 1.76 (0.85–3.62) | 0.13 |
| M2 (other +/- lung met; *n*=30) | 4.86 (2.65–8.91) | <0.01 |
| SOX2<br>high (>10%) | 3.19 (1.74–5.84) | <0.01 |
| Age<br>≥15 years | 1.34 (0.79–2.27) | 0.28 |
| Site<br>axial | 1.64 (0.90–2.96) | 0.11 |

To exclude interdependency of metastasis and SOX2 expression status, we repeated the multivariate analysis considering only patients with localized disease (M0, *n*=87). In this analysis, SOX2-high expression (HR=3.22, *P*<0.01) represented the main risk-factor for survival, followed by age (≥15 years, HR=2.43, *P*=0.04), whereas primary axial tumor site (HR=1.66, *P*=0.25) showed only a tendency to affect survival (**Table 3**; *n*=87). In agreement with these findings, in both validation- and test-cohorts, univariate analysis did not show a correlation between SOX2-high and metastasis (**Table 4**, *n*= 141, *P*=0.8; and **Table 5**, *n*=76, *P*=0.5). Taken together, these findings demonstrate that SOX2-high expression constitutes an independent risk-factor for survival of EwS patients, even in patients with localized disease.

**Table 3.**
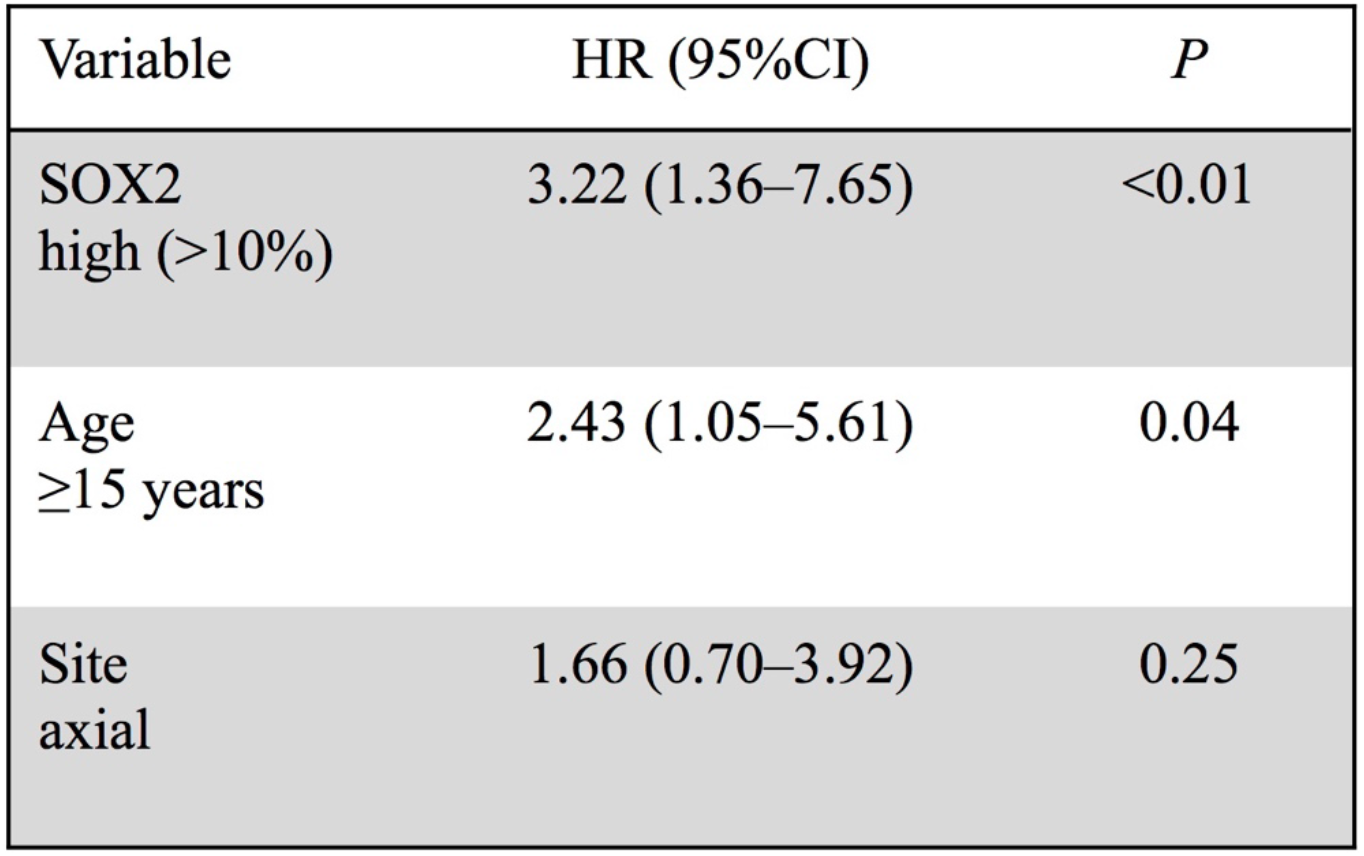
Multivariate analysis in patients with localized disease from the TMA-cohort (*n*=87).

### SOX2-high expression significantly correlates with tumor relapse

Next, we investigated whether the correlation between SOX2-high expression and OS is associated with histological response, high tumor volume (≥200 ml), and relapse, which are risk-factors for worse outcome^12,22–24^.

In our TMA-cohort, data on histological response, tumor volume and tumor relapse were available for 84, 136 and 141 patients, respectively (**Table 4**). Our data did not show an association of SOX2-high expression with histological response (**Table 4**). However, we observed a tendency for an association between SOX2-high and tumor volume. In fact, 58.3% of patients with high SOX2 expression had a tumor volume ≥200 ml compared to 41.7% patients with low SOX2 expression (*P*=0.23) (**Table 4**). Since the association of SOX2 and higher tumor volume suggested a role for SOX2 in EwS growth, we performed functional *in vitro* and *in vivo* experiments using an established EwS cell line model (POE) for which we generated a derivate with a doxycycline (dox-)inducible shRNA against *SOX2*. The results showed that knockdown of *SOX2* in POE cells led to strong reduction in cell proliferation, clonogenic growth and anchorage-independent growth compared to cells expressing a nontargeting control shRNA (**Fig. S2**). Consistently, dox-induced silencing of SOX2 in POE cells xenografted in immunocompromised NSG mice significantly reduced tumor growth (**Fig. S3**). In addition, we noted a striking correlation of SOX2-high expression with tumor relapse (*P*=0.001) (**Table 4**). While only 38.7% of patients with SOX2-low expression had relapse, recurrence occurred in 77.3% of patients with SOX2-high expression (**Table 4**), which may explain the observed poor outcome for these patients. In agreement with these observations, gene set enrichment analysis (GSEA) in our mRNA cohort showed that genes co-expressed with *SOX2* in primary EwS tumors are involved in stemness, proliferation, dedifferentiation and cancer relapse (**Fig. S2A, Supplementary Table 1**).

**Table 4.** Correlation of SOX2 immunoreactivity with metastasis, histological response, tumor volume and relapse in the TMA-cohort (*n*=141).

|  | Metastasis at diagnosis<br>(n=141) |  |  | Histological response<br>(n=84) |  |  | Tumor volume<br>(n=136) |  |  | Relapse<br>(n=141) |  |  |
| --- | --- | --- | --- | --- | --- | --- | --- | --- | --- | --- | --- | --- |
|  | No | Yes | P | Good | Poor | P | <200 ml | ≥200 ml | P | No | Yes | P |
| SOX2 |  |  |  |  |  |  |  |  |  |  |  |  |
| low | 74 (62.2%) | 45 (37.1%) | 0.8 | 57 (79.2%) | 15 (20.8%) | 1.0 | 67 (58.3%) | 48 (41.7%) | 0.23 | 73 (61.3%) | 46 (38.7%) | 0.001 |
| high | 13 (59.1%) | 9 (40.1%) |  | 10 (83.3%) | 2 (16.7%) |  | 9 (42.9%) | 12 (57.1%) |  | 5 (22.7%) | 17 (77.3%) |  |

**Table 5.**
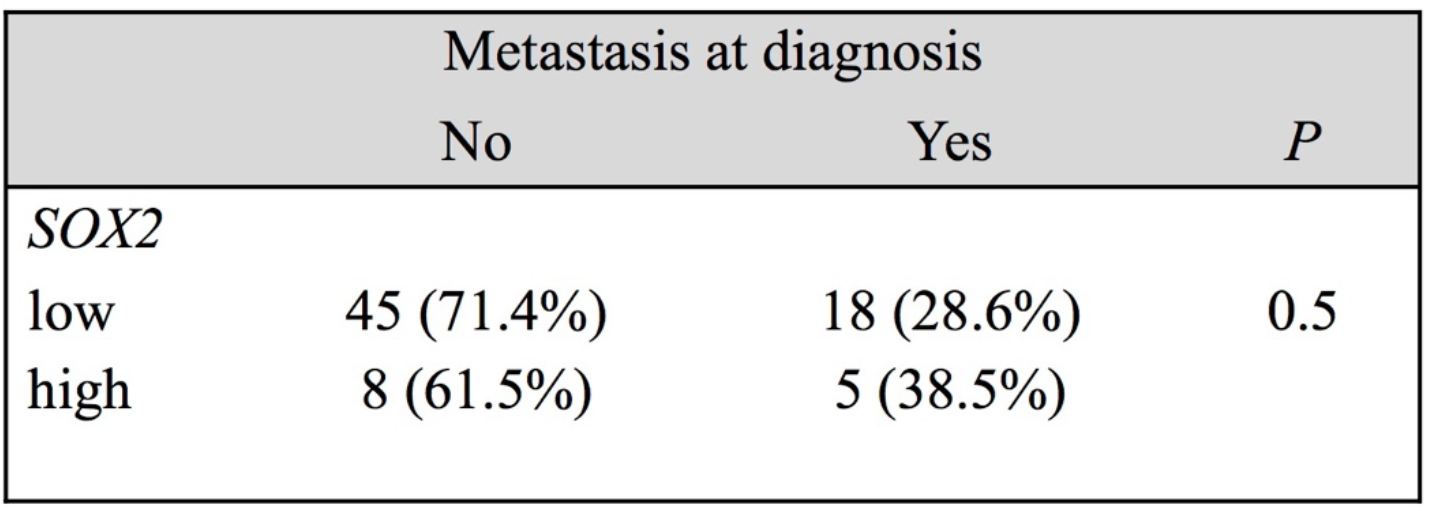
Correlation of *SOX2* mRNA with metastasis in the mRNA-cohort (*n*=76). Information on the status of metastasis was only available for a subset of the test cohort.

|  | Metastasis at diagnosis |  | P |
| --- | --- | --- | --- |
|  | No | Yes |  |
| SOX2 |  |  |  |
| low | 45 (71.4%) | 18 (28.6%) | 0.5 |
| high | 8 (61.5%) | 5 (38.5%) |  |

Collectively, these results suggest that SOX2-high expression may confer worse outcome to EwS patients by contributing to tumor growth and relapse.

## DISCUSSION

Up to 30–40% of Ewing sarcoma (EwS) patients with non-metastatic disease develop local or metastatic relapse within a time range of 2–10 years^25^. Clinical prognostic markers such as primary dissemination, tumor site, size, age and histological response to chemotherapy are established risk-factors used for therapeutic stratification^25^. However, especially for patients with localized disease and otherwise no apparent risk-factors, risk prediction is challenging^26^. Although there is broad consensus that clinical management will benefit from prognostic or predictive biomarkers that can guide therapeutic decisions, there are currently no *bona fide* biomarkers available that may help predicting tumor relapse and outcome of patients with EwS^1,12^.

In this study, we report, for the first time, that high mRNA or protein expression of SOX2 is a stratification risk-factor for ~16–20% of EwS patients with poor outcome. Notably, multivariate analyses including either all or only patients with localized disease demonstrated that SOX2-high expression represents an independent and strong risk-factor for EwS patients. In line with our findings in EwS, high expression or amplification of *SOX2* has been reported to correlate with poor survival in breast, colorectal, esophageal, laryngeal, endometrial and ovarian carcinoma^14,27–30^. Similarly to EwS, SOX2 is highly expressed in a subset of patients with lung adenocarcinoma and was found to be an independent predictor of poor survival^31^.

Prior reports suggested that *SOX2* may constitute a direct EWSR1-FLI1 target gene^13,32^. Surprisingly, we found that the vast majority of molecularly confirmed EwS cases do neither express SOX2 on the mRNA nor on the protein level, suggesting that the mode of regulation and the functional role of SOX2 may be more complex than anticipated.

In our TMA-cohort, SOX2-high expression showed a correlation with tumor relapse (*P*=0.001), which likely explains the poor outcome of these patients. In support of our findings, it has been shown in sinonasal carcinomas that patients with *SOX2* amplification had a significantly higher rate of tumor recurrences than those without *SOX2* amplification^33^. However, in our TMA-cohort 38.7% of patients with SOX2-low expression showed relapse suggesting that in this subset of patients a different driver might mediate tumor relapse. Although the correlation between SOX2-high and tumor volume was not significant in our TMA-cohort, in line with previous reports in EwS^13,34^, our *in vitro* and *in vivo* results confirmed a functional role of SOX2 in EwS cell proliferation and tumor growth suggesting that a larger cohort might enable validation of this correlation in EwS patients. In agreement, our GSEA in primary EwS indicated that SOX2-high tumors are enriched for gene signatures involved in dedifferentiation, proliferation and stemness, which may promote tumor relapse. Interestingly, GSEA also revealed that *SOX2* co-expressed genes overlap with genes involved in relapse of malignant melanoma – a tumor of neuroectodermal origin^35^, which is also proposed for EwS^36^. In support of these data, GSEA of differentially expressed genes upon *SOX2* knockdown in three SOX2-high EwS cell lines demonstrated a significant upregulation of gene signatures involved in differentiation-related processes such as neurite outgrowth, integrin cell surface interaction, and components of the basement membrane (**Fig. S1B, Supplementary Table 2**).

Despite the exploratory nature of this study, SOX2-high expression may constitute the first promising biomarker for outcome prediction and stratification of high-risk EwS patients with localized disease, which is readily available due to standardized assessment by qRT-PCR and/or IHC. Moreover, our results suggest that SOX2-high EwS patients may benefit from an alternative therapy that could be administered upfront. Therefore, we recommend validation of these observations in additional prospective studies and to experimentally elucidate the precise molecular role of SOX2 in EwS.

## ACKNOWLEDGEMENTS

The laboratory of T. G. P. Grünewald is supported by grants from the ‘Verein zur Förderung von Wissenschaft und Forschung an der Medizinischen Fakultät der LMU München (WiFoMed)’, by LMU Munich’s Institutional Strategy LMUexcellent within the framework of the German Excellence Initiative, the ‘Mehr LEBEN für krebskranke Kinder – Bettina-Bräu-Stiftung’, the Walter Schulz Foundation, the Wilhelm Sander-Foundation (2016.167.1), the Friedrich-Baur foundation, the Matthias-Lackas foundation, the Barbara & Hubertus Trettner foundation, the Dr. Leopold und Carmen Ellinger foundation, the Gert & Susanna Mayer foundation, the Deutsche Forschungsgemeinschaft (DFG 391665916), and by the German Cancer Aid (DKH-111886 and DKH-70112257). J. Li was supported by a scholarship of the China Scholarship Council (CSC), J. Musa was supported by a scholarship of the Kind-Philipp foundation, and T. L. B. Hölting by a scholarship of the German Cancer Aid. M. F. Orth and M. M. L. Knott were supported by scholarships of the German National Academic Foundation. G. Sannino was supported from a scholarship from the Fritz-Thyssen Foundation (FTF-40.15.0.030MN). The work of U. Dirksen is supported by grants from the German Cancer Aid (DKH-108128 and DKH-70112018), the ERA-Net-TRANSCAN consortium (project number 01KT1310), and Euro Ewing Consortium (EEC, project number EU-FP7 602856), both funded under the European Commission Seventh Framework Program FP7-HEALTH (http://cordis.europa.eu/), the Barbara & Hubertus Trettner foundation, and the Gert & Susanna Mayer foundation. G. Hardiman was supported by grants from the National Science Foundation (SC EPSCoR) and National Institutes of Health (U01-DA045300). The authors thank A. Sendelhofert and A. Heier for excellent technical support.

## AUTHOR CONTRIBUTIONS

**Conception and design:** Giuseppina Sannino and Thomas G. P. Grünewald.

**Provision of study material and patients:** Uta Dirksen, Heribert Jürgens, Wolfgang Hartmann, Thomas Kirchner, Ana Sastre, Javier Alonso.

**Financial and administrative support:** Thomas Kirchner and Thomas G. P. Grünewald.

**Data analysis and interpretation:** Giuseppina Sannino, Andreas Ranft, Susanne Jabar, Uta Dirksen, Aruna Marchetto, Martin F. Orth, Julia S. Gerke, Willian De Silveira and Gary Hardiman, Fabienne S. Wehweck, Merve M. Kiran, Thomas G. P. Grünewald.

**Experimental support:** Constanze Zacherl, Rebeca Alba-Rubio, Stefanie Stein, Tilman L. B. Hölting, Julian Musa, Laura Romero-Pérez, Florencia Cidre-Aranaz, Maximilian M. L. Knott, Jing Li.

**Manuscript writing:** Giuseppina Sannino, Uta Dirksen and Thomas G. P. Grünewald.

**Final approval of the manuscript:** All the authors.

